# Dynamic saccade context triggers more stable object-location binding

**DOI:** 10.1101/2023.04.26.538469

**Authors:** Zitong Lu, Julie D. Golomb

**Author notes:** Please address correspondence to: Julie D. Golomb The Ohio State University Department of Psychology 1835 Neil Avenue Columbus, OH, 43210.

## Abstract

Our visual systems rapidly perceive and integrate information about object identities and locations. There is long-standing debate about how we achieve world-centered (spatiotopic) object representations across eye movements, with many studies reporting persistent retinotopic (eye-centered) effects even for higher-level object-location binding. But these studies are generally conducted in fairly static experimental contexts. Might spatiotopic object-location binding only emerge in more dynamic saccade contexts? In the present study, we investigated this using the Spatial Congruency Bias paradigm in healthy adults. In the static (single saccade) context, we found purely retinotopic binding, as before. However, robust spatiotopic binding emerged in the dynamic (multiple frequent saccades) context. We further isolated specific factors that modulate retinotopic and spatiotopic binding. Our results provide strong evidence that dynamic saccade context can trigger more stable object-location binding in ecologically-relevant spatiotopic coordinates, perhaps via a more flexible brain state which accommodates improved visual stability in the dynamic world.

**Significance Statement:** One of the most fundamental challenges for human behavior is how we integrate and stabilize perceptual information in our ever-changing sensory environments. In particular, we make multiple eye movements every second, constantly displacing and distorting our visual input. Yet despite receiving visual input in these disjointed, eye-centered (retinotopic) coordinates, we perceive the world as stable, based on objects’ world-centered (spatiotopic) locations. Our study provides strong evidence for a previously unstudied cue – dynamic saccade context – in triggering more stable object-location binding, which offers a novel step forward in understanding how we form a stable perception of the dynamic world. More broadly, these findings suggest the importance of considering dynamic saccade context in visual perception and cognitive neuroscience studies.

## Introduction

One of the most fundamental challenges for human behavior is how we integrate and stabilize perceptual information in our ever-changing sensory environments. In particular, we make multiple eye movements every second, constantly displacing and distorting our visual input. Understanding how the human brain attains stable visual perception requires understanding both how spatial information is stabilized across eye movements and how this spatial information is integrated with visual feature and object representations.

As a real-world example, when we are searching for a red pen on a desk, we are able to not only recognize the shape and color of the pen but also the location of the pen. A great deal of research has gone into understanding how information about object identity and location are combined, often called object-location binding (Treisman, 1996). Traditionally, information regarding the ‘where’ and ‘what’ of an object has been considered to be processed through separate cognitive and neural processes (Goodale & Milner, 1992; Mishkin & Ungerleider, 1982). However, increasing behavioral and neuroimaging studies have found that spatial location interactions can be automatically encoded and bound to an object’s representation during object recognition (Chen, 2009; Cichy et al., 2011; Golomb et al., 2014; Kovacs & Harris, 2019; Schwarzlose et al., 2008; Treisman & Gelade, 1980; Treisman & Zhang, 2006; Tsal & Lavie, 1988).

However, as we view the world and move our eyes, the input location of objects onto our eyes is constantly changing. This prompts the question, what is the spatial reference frame for object-location binding? When searching for the pen on the desk, we perceive the pen as having a static position in the world (spatiotopic, world-centered coordinates), even when eye movements change the pen’s location on the retina (retinotopic, eye-centered coordinates). Thus, our brains should have the ability to link together an object’s identity and its spatiotopic location when we are viewing something in the real world. However, evidence for spatiotopic object-location binding has proved elusive. Even outside the context of object-location binding, it is controversial whether and how the brain represents spatial information in spatiotopic coordinates (Duhamel et al., 1992, 1997; Gardner et al., 2008; Golomb et al., 2008; Golomb & Kanwisher, 2012; Marino & Mazer, 2016; Melcher & Morrone, 2003; Snyder et al., 1998; Turi & Burr, 2012). It is still unresolved how object features and identity are processed across eye movements, which has been referred to as the ‘hard binding problem’ (Cavanagh et al., 2010; Golomb & Mazer, 2021).

One recent behavioral paradigm, Spatial Congruency Bias (Golomb et al., 2014), provides a robust measure of object-location binding and the spatial reference frame of binding (Shafer-Skelton et al., 2017). In the standard Spatial Congruency Bias task with no saccade, participants are asked to judge whether two objects presented sequentially were of the same identity or not. Although object location is irrelevant to the task, if the two sequential objects appeared in the same location, participants are more likely to judge them as the same identity, in contrast to if they appeared in different locations, showing that object location is automatically bound to and fundamentally influences perception of object identity. To investigate object-location binding across eye movements, Shafer-Skelton et al., 2017 added a saccade during the delay between objects to distinguish retinotopic and spatiotopic bindings. Strikingly, they found that the Spatial Congruency Bias was preserved across the saccade, but entirely based in retinotopic coordinates, with no evidence for spatiotopic object-location binding even at longer post-saccade delays or for complex objects requiring higher-level processing (Shafer-Skelton et al., 2017).

Why don’t we find spatiotopic object-location binding across saccades, when that is clearly the more ecologically relevant coordinate system for behavior? One option is that despite ecological relevance, visual information is simply always coded in native retinotopic coordinates. But an alternative option is that spatiotopic object-location binding might emerge only under certain contexts, and the prior studies were not designed in a way to tap into this. For example, Shafer-Skelton et al., 2017 study and other studies referenced above tend to rely on fairly static contexts to explore reference frames, where participants are asked to fixate on one location for an extended period of time, and perhaps execute a single saccade on each trial. In contrast, in real life we typically execute multiple eye movements in rapid succession. Some models propose different spatiotopic localization mechanisms relying on active versus passive feedback (Bergelt & Hamker, 2019; Golomb et al., 2011; Ross & Ma-Wyatt, 2003; Sun & Goldberg, 2016; Wexler & Van Boxtel, 2005), perhaps accommodating more tolerant or optimally updated spatial representations in the case of multiple eye movements (Poletti et al., 2013; Zhang et al., 2020).

Our postulation is that dynamic contexts with multiple active eye movements may facilitate a more stable and integrative state, which could be particularly valuable during object recognition, and thus offer an important clue for how our visual systems solve the ‘hard binding problem’. In other words, eye movements may be thought of as not only part of the challenge of visual stability, but also part of the solution. We hypothesize that our visual system has the ability to bind object information to spatiotopic coordinates under more dynamic saccade contexts, which are more ecological. Perhaps previous studies only found retinotopic object-location binding because they did not fully induce the spatiotopic binding mechanism. In the present study, our goal is to revisit the behavioral object-location binding question and ask whether dynamic saccade context, in contrast to more static contexts, can trigger spatiotopic object-location binding.

## Method

### Overview

In Experiment 1, we tested a dynamic saccade context where participants performed continual eye movements during the task and were asked to judge if two objects presented sequentially were the same identity or not at the end of each trial. In comparison, Experiment 2 was a control version of the task testing a static condition where participant were asked to conduct only a single saccade during the delay between the two stimuli. We hypothesized that if the dynamic saccade context could trigger spatiotopic object-location binding, we might see a spatiotopic bias in Experiment 1; otherwise, we would expect similar retinotopic-only results in Experiment 1 and Experiment 2. We then conducted two additional experiments (Experiment 3 and Experiment 4) to further isolate specific factors that might contribute to the influence of dynamic saccade context on spatiotopic object-location binding.

### Subjects

The research was approved by the Ohio State University Behavioral and Social Sciences Institutional Review Board. Each of the four experiments included 16 subjects (with a different set of subjects in each experiment). All subjects reported normal or corrected-to-normal vision and gave informed consent. Subjects were compensated with course credit or payment. Sample size was chosen in advance based on power analyses of previous spatial congruency bias studies. A power analysis of the original spatial congruency bias effect (Experiment 1 of Golomb et al., (Golomb et al., 2014)), which had an effect size of dz = 1.01 for the comparison of SameLocation versus DifferentLocation bias, estimated *N*=13 would be needed to achieve.9 power. Also, a power analysis of previous spatial congruency bias effect with one saccade (Experiment 4 of Shafer-Skelton et al (Shafer-Skelton et al., 2017)), which had an effect size of dz = 0.92 for the comparision of RetinotopicLocation and ControlLocation, estimated *N* = 15 would be needed to achieve.9 power. We set sample size at N = 16 (matching prior studies), which should have sufficient power. Additional subjects completed the experiment but were excluded due to poor task performance (overall accuracy < 55% or hit rate < 50% or false alarm rate > 50 %; predetermined threshold).

### Experimental setup

Stimuli were presented using Psychtoolbox extension (Brainard, 1997) for MATLAB (Math Works), on a 21-in (53.34-cm) flat screen CRT monitor. Subjects were seated at a chinrest 60 cm from the monitor.

### Eye tracking

Eye position was monitored with an EyeLink 1000 eye-tracking system recording pupil and corneal reflection position. Fixation was monitored for all experiments.

### Stimuli

Stimuli were the same as those in Golomb et al., (Golomb et al., 2014), from the Tarr stimulus set (stimulus images courtesy of Michael J. Tarr, Center for the Neural Basis of Cognition and Department of Psychology, Carnegie Mellon University, http://www.tarrlab.org). Stimuli were drawn from ten families of shape morphs; within each family, the “body” of the shape remained constant, while the “appendages” could vary in shape, length, or relative location. The Stimulus 1 shape was randomly chosen for each trial. On Same Shape trials, the Stimulus 2 shape was an identical image. On Different Shape trials, the second shape was chosen as a different shape from the same morph family. We used the easiest morph level (the two images with the greatest morph distance within a family) for all subjects instead of individually staircasing task difficulty because in Golomb et al., (Golomb et al., 2014) and Shafer-Skelton et al (Shafer-Skelton et al., 2017), participants were already within the desired accuracy range at this easiest morph level (maximum staircase value) in both no-saccade and saccade tasks. Stimuli were sized 6.25°×6.25°, and stimulus orientation was never varied.

### General procedure

All experiments used the same stimuli, and subjects needed to follow the fixation point throughout the task. There were four possible fixation locations, centered on the screen and forming the corners of an invisible 10°×10° square, and nine possible stimulus locations, forming a 3×3 grid such that each fixation location had four adjacent stimulus locations of equal eccentricity with 7.06°. Before the main task, subjects were asked to do a saccade pre-task and a practice task. For the saccade task, the location of fixation changed (from 4 possible fixation locations, only vertical or horizontal change) 50 times (once per second), and subject followed the fixation changes to do 50 saccades. We calculated the mean saccade time for each subject as the individual saccade reaction time used in the main task. For the practice task, each subject completed eight trials which were consistent with the trial in the main task.

### Experiment 1: Dynamic saccade context

Figure 1A shows a sample trial timeline for Experiment 1. On each trial, the fixation cross first jumped back and forth between two Fixations (Fixation 1 and Fixation 2). During this first part of the trial, Stimulus 1 was presented. Then there was a critical saccade in an orthogonal direction to Fixation 3. Then the fixation cross jumped back and forth between Fixations 3 and 4; during this second part of the trial, Stimulus 2 was presented. The fixation cross stayed at each location for 1000ms. The fixation cross started at Fixation 1, and it changed 5 times between Fixation 1 and Fixation 2. Then the fixation cross jumped from Fixation 2 to Fixation 3, then changed twice between Fixation 3 and Fixation 4. Subjects were asked to follow the fixation cross throughout the task (monitored via eye-tracking). There were 8 possible eye movement routes (Figure 1D).

**Figure 1.**
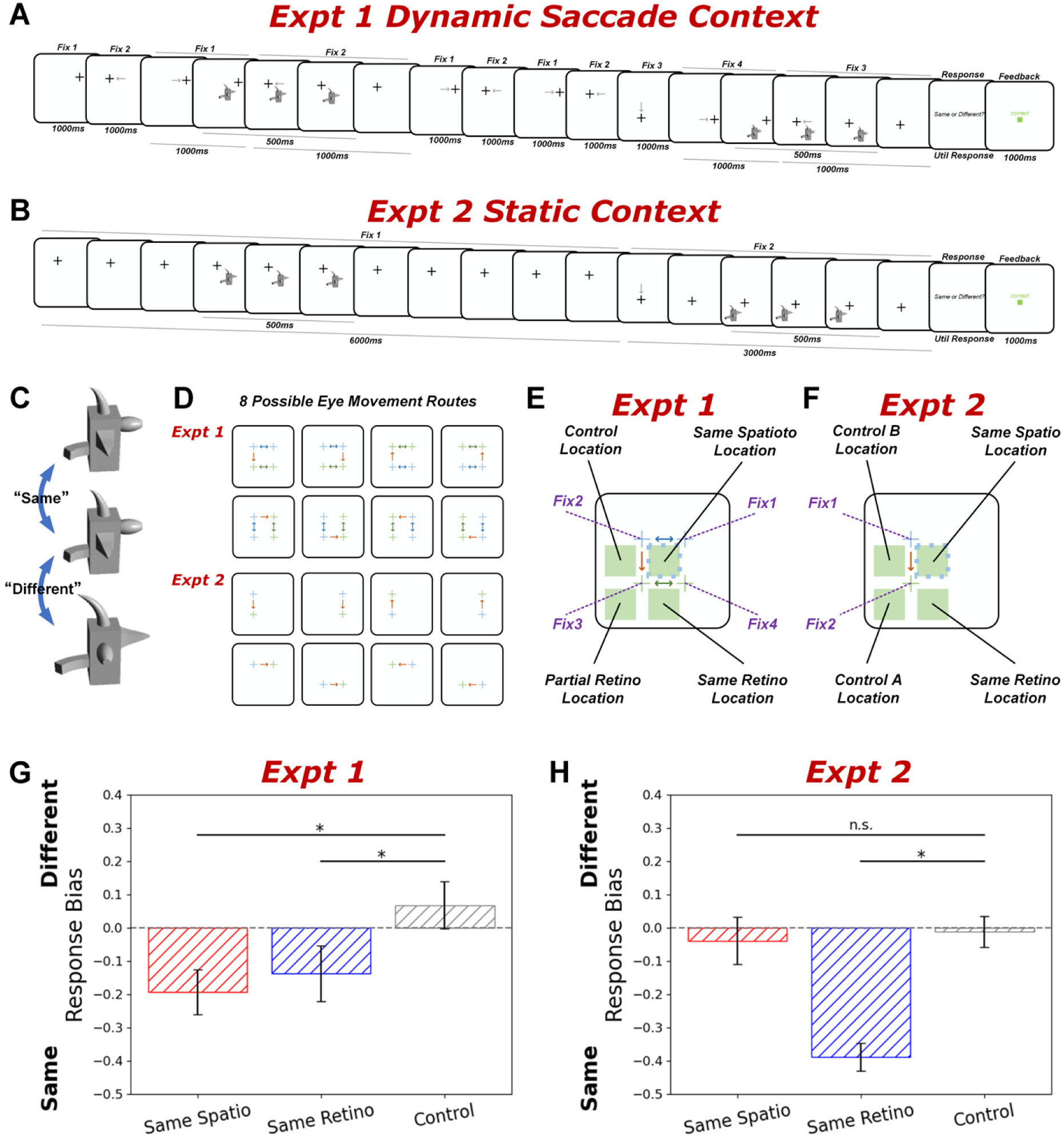
Methods and results of Experiment 1 and 2. (A) Trial timing for Experiment 1. While doing dynamic saccades, subjects saw two sequential object presentations. Subjects completed 8 saccades on each trial, two of which happened during the time the two stimuli appeared. The task was to judge whether two stimuli were the same or different identity. (B) Trial timing for Experiment 2. Subjects saw two sequential object presentations on each trial. Subjects completed only 1 saccade during the delay between two stimuli on each trial. The task was to judge whether two stimuli were the same or different identity. (C) Sample identity stimuli from a certain shape morph family. Left two stimuli are different, and right two stimuli are same. (D) 8 possible eye movement routes in two experiments. The brown arrow in Expt 1 was a critical saccade in an orthogonal direction to Fixation 3, which was consistent with the only arrow in Expt 2. (E) An illustration of 4 possible location conditions for Experiment 1. For a given Stimulus 1 location (blue dotted square), there were four possible locations for Stimulus 2 (indicated by four squares): Same Spatiotopic Location (same screen position as Stimulus 1), Same Retinotopic Location (same location as Stimulus 1 relative to 2 fixations during stimulus), Partial Retinotopic Location, and Control Location (different spatiotopic and retinotopic location). (F) An illustration of 4 possible location conditions for Experiment 2. For a given Stimulus 1 location (yellow dotted square), there were four possible locations for Stimulus 2 (indicated by four squares): Same Spatiotopic Location (same screen position as Stimulus 1), Same Retinotopic Location (same location as Stimulus 1 relative to the fixation), Control A Location (different spatiotopic and retinotopic location), and Control B Location (different spatiotopic and retinotopic location). (G) Response bias (criterion) on the identity task plotted for Same Spatiotopic, Same Retinotopic, and Control location conditions, for Experiment 1. A more negative response bias indicates a greater tendency to respond ‘Same Identity’. (H) Response bias (criterion) on the identity task plotted Same Spatiotopic, Same Retinotopic, and Control (the average of Control A and Control B) location conditions, for Experiment 2. Error bars are standard error of the mean. Asterisk indicates p<.05 (paired t-tests between different conditions). Spatio = Spatiotopic; Retino = Retinotopic. See Supplemental Materials for tables with full results for all behavioral measures and conditions.

Each object stimulus was shown for 500ms, and was designed to straddle an eye movement, such that the eye movement would occur during the stimulus presentation for full dynamic context. Ideally Stimulus 1 would be shown from approximately 250ms before to 250ms after the 3rd saccade, with Stimulus 2 straddling the 8th saccade with similar timing. Because there is variability in saccadic reaction times across individuals and trials, we did the following: First, we estimated each individual’s average saccade reaction time (aSRT) from the saccade pre-task. Then, in the main task, we used the individualized aSRT to determine the onset of the stimulus relative to the saccade cue. For example, if saccade 3 was cued at t=3000ms, and that subject’s aSRT was 185ms, then Stimulus 1 would be presented at 2935ms (saccade cue onset + aSRT - 250). We used an eye-tracker to monitor eye position to make sure the subjects made the saccade while the stimulus was presented. Moreover, if at any point the subject’s fixation deviated greater than 2° from the current fixation, the trial was aborted and repeated later in the block.

After the last fixation, subjects saw the question ‘Same or Different’ and were instructed to make a two-alternative forced choice same/different judgement comparing the two objects’ identities (shapes: Figure 1C); location was irrelevant to the task. Subjects responded by button press (‘j’ for ‘Same’ and ‘k’ for ‘Different’) and were presented with visual feedback (‘correct’ or ‘incorrect’ in green or red on the screen). They were also provided feedback if they broke fixation at any time during the trial (‘Please do the saccade faster!’ on the screen) or didn’t give a response within 3 seconds (‘No response’ on the screen).

Stimulus 2 could appear in one of four possible location conditions (25% trials for each) (Figure 1E). The two main conditions of interest were Same Spatiotopic Location (the same absolute screen location as Stimulus 1) and Same Retinotopic Location (the same location as Stimulus 1 relative to 2 fixations during stimulus). These were compared to the Control Location condition (different spatiotopic and retinotopic location but at an equal eccentricity from Fixation 2). Finally, because the stimulus straddled the saccade and could have stimulated two different retinotopic positions, we also included a Partial Retinotopic Location condition (the same retinotopic location as Stimulus 1 relative to only one of 2 fixations during stimulus). We did not have clear hypotheses for this condition and do not analyze it further, but it was important to include as a separate condition so as not to contaminate either the Control Location or Same Retinotopic Location conditions.

The location of Stimulus 1 was chosen from one of two possible locations for a given eye movement route (based on the saccade from Fixation 2 to Fixation 3, such that all four Stimulus 2 location conditions were possible). For example, if Fixation 1 and Fixation 2 were the upper-right and upper-left fixation positions and the saccade from Fixation 2 to Fixation 3 was downward, Stimulus 1 could appear in either the middle-left or middle-middle position on the screen, such that the Same Spatiotopic, Same Retinotopic, Partial Retinotopic, and Control locations of Stimulus 2 were located at equal eccentricity from Fixation 3, which was the bottom-left fixation position. The identity of Stimulus 1 was chosen randomly, and Stimulus 2 could have either the same or different identity as Stimulus 1. These sixteen conditions (two Stimulus 1 location conditions × four Stimulus 2 location conditions × two identity conditions), along with eight possible eye movement routes, were counterbalanced and equally likely.

Subjects completed 8 blocks and 32 trials per block (256 trials in total, two trials for each of the 128 Stimulus 1 locations × Stimulus 2 location × identity × eye movement route conditions, in randomized order and randomly divided into 8 blocks), in addition to any trials that were aborted due to eye-tracking errors (which were repeated later in a randomized order within the same block).

### Experiment 2: Static context

Figure 1B shows a sample trial timeline for Experiment 2. Unlike Experiment 1, Experiment 2 only had the one critical saccade in the middle of the trial. For a trial, the fixation cross began at one fixation location (Fixation 1) and remained there for 6000ms (matching the overall timing of Experiment 1). Stimulus 1 appeared for 500ms during this period, with the same timing as Experiment 1. Then, the fixation cross jumped to either the adjacent vertical or horizontal fixation location (Fixation 2) and stayed at the new fixation location for 3000ms. Stimulus 2 was presented for 500ms during this second period, again with the same timing in Experiment 1. There were 8 possible eye movement routes (Figure 1F).

Stimulus 2 could appear in one of four possible location conditions (25% trials for each) (Figure 1F). The two main conditions of interest were again the same absolute screen location (Same Spatiotopic Location), and the same location as Stimulus 1 relative to fixation (Same Retinotopic Location). Because there was never a saccade during the stimulus, there was no Partial Retinotopic condition in Experiment 2, so there were two control locations at an equal eccentricity from Fixation 2 (Control A Location and Control B Location). All other details were the same as Experiment 1. All conditions were counterbalanced and equally likely. Subjects completed 8 blocks with 32 trials per block, in addition to any trials that were aborted due to eye-tracking errors.

### Experiment 3: Partial dynamic context with repeated eye movements only

The task was modified from Experiment 1. The timing of each trial was the same as in Experiment 1, except the onset times of the two stimuli were shifted such that they appeared during a fixation period instead of straddling a saccade (Figure 2A). In Experiment 3, ideally Stimulus 1 was shown from approximately 250ms to 750ms after the 3^rd^ saccade completion time, and stimulus 2 from approximately 250ms to 750ms after the 8^th^ saccade completion time. Like Experiment 1, because there was variability in saccadic reaction times across individuals and trials, we used the individualized aSRT to determine the saccade completion time and the onset of the stimulus. For example, if saccade 3 was cued at t=3000ms, and that subject’s aSRT was 185ms, then Stimulus 1 would be presented at 3435ms (saccade cue onset + aSRT + 250). We utilized an eye-tracker to make sure that subjects were looking at only one fixation point when a stimulus appeared.

**Figure 2.**
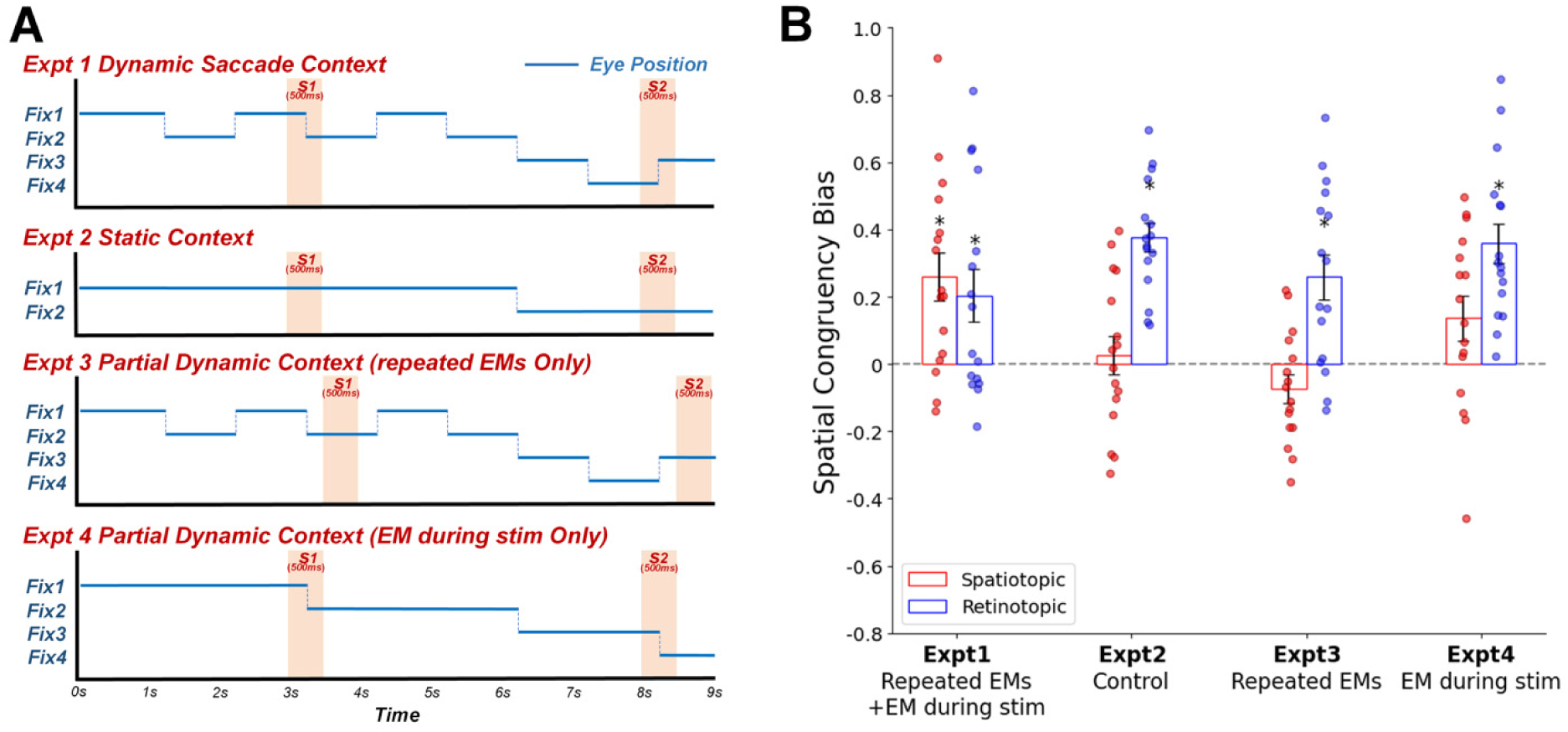
(A) Trial timing for all four experiments. The X-axis represents the time for each trial. The Y-axis represents the four possible fixation locations. The blue lines indicate eye positions during the task. The orange blocks indicate two stimuli on each trial. (B) Spatial congruency bias on the identity task plotted for spatiotopic and retinotopic conditions (spatiotopic congruency bias: Control bias minus Same Spatiotopic bias; retinotopic congruency bias: Control bias minus Same Retinotopic bias). Error bars are standard error of the mean. Asterisk indicates p<.05 (one-sample t-test for each condition). Spatio = Spatiotopic; Retino = Retinotopic; EM = eye movement, stim = stimulus. See Supplemental Materials for tables with full results for all behavioral measures and conditions.

Stimulus 2 could appear in four possible locations, Same Spatiotopic Location, Same Retinotopic Location, Control A Location and Control B Location, similar to Experiment 2. All conditions were counterbalanced and equally likely. Subjects completed 8 blocks and 32 trials per, in addition to any trials that were aborted due to eye-tracking errors.

### Experiment 4: Partial dynamic context with eye movement during stimulus only

The task was also modified from Experiment 1. In Experiment 4 the timing of the stimuli was the same as in Experiment 1 (eye movement during stimulus), but now there were fewer saccades. In order to preserve the design, we needed a minimum of three saccades per trial: One saccade during each stimulus, and the critical saccade in the middle of the trial to distinguish spatiotopic and retinotopic locations (Figure 2A). In Experiment 1, whereas subjects made four saccades between Fixation 1 and Fixation 2 in the first part of the trial; in Experiment 4 subjects only made one saccade between Fixation 1 and Fixation 2. To best match overall timing, this first saccade occurred with the same timing as the 3^rd^ saccade in Experiment 1, during the presentation of Stimulus 1. Similarly, subjects only made one saccade in the second part of the trial between Fixation 3 and Fixation 4, with the same timing as the 8^th^ saccade in Experiment 1, during the presentation of Stimulus 2. The critical (middle) saccade was cued at the same time in all four experiments (Figure 2A).

Stimulus 2 could appear in four possible locations, the same as Experiment 1: Same Spatiotopic Location, Same Retinotopic Location, Partial Retinotopic Location and Control Location, similar to Experiment 1. All conditions were counterbalanced and equally likely. Subjects completed 8 blocks and 32 trials per block, in addition to any trials that were aborted due to eye-tracking errors.

### Analysis

Our primary measure for all experiments was the spatial congruency bias, which is calculated as the difference in response bias for the same location versus a different location condition (Golomb et al., 2014). For each subject, we calculated hit and false alarm rates for each of the four location conditions. We defined a ‘hit’ as a ‘Same’ response when the two stimuli were actually the same (Same Identity condition), and a ‘false alarm’ as a ‘Same’ response when the two stimuli were different (Different Identity condition). Using signal detection theory, we applied the standard formula (Stanislaw & Todorov, 1999) to calculate response bias (criterion) for each subject, for each location condition:

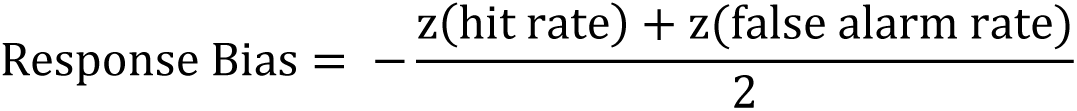

Note that, although it is often assumed that response bias measures reflect decision-level effects, response bias can also reflect perceptual-level processes, as in the case of the spatial congruency bias (Babu et al., 2023; Golomb et al., 2014; Shafer-Skelton et al., 2017; Witt et al., 2015).

To evaluate the spatial congruency bias (i.e., whether there was a greater bias to report two stimuli as the same identity when they appeared in the same retinotopic and//or spatiotopic location compared to a different location), we compared the response bias for the control location conditions to the response bias for the same spatiotopic and same retinotopic location conditions. For Experiments 1 and 4, the single “different” location (Control) was used for both Spatiotopic and Retinotopic comparisons; For Experiments 2 and 3, we used the average of Control A and Control B as the “different” location (Control) for both Spatiotopic and Retinotopic comparisons. We calculated the spatiotopic spatial congruency by subtracting Same Spatiotopic bias from Control bias, and the retinotopic spatial congruency bias by subtracting Same Retinotopic bias from Control bias. (Full comparisons across all pairs of conditions can also be found in Tables S3, S5, S7, and S9.)

The spatial congruency biases were calculated separately for each subject. One-sample t-tests were used to determine whether spatiotopic and retinotopic spatial congruency biases were significantly different from zero. Paired t-tests were used to determine whether spatiotopic and retinotopic congruency biases were significantly different from each other. We also report effect sizes using Cohen’s d, and calculated Bayes factors of both one-sample and paired t-tests. We report both the frequentist and Bayesian statistics here.

To compare the effects between dynamic saccade and static fixation contexts, we performed a 2 × 2 ANOVA on the first two experiments with a within-subjects factor of Reference frame (Spatiotopic or Retinotopic) and an across-subjects factor of Experiment Context factor (dynamic saccade or static fixation) for spatial congruency bias.

In addition, we conducted an across-experiments analysis across all four experiments to further explore how the two dynamic context factors (repeated eye movements and eye movements during stimulus) influence object-location binding. First, we performed a 2 × 2 × 2 ANOVA with a within-subjects factor of Reference frame (Spatiotopic or Retinotopic) and two across-subjects factors of Repeated eye movements (had or not) and Eye movement during stimulus (had or not) for spatial congruency bias. Then, we followed with two 2 (Repeated eye movements factor) × 2 (Eye movement during stimulus factor) ANOVAs to assess if there was a significant main effect or interaction for Spatiotopic and Retinotopic congruency biases respectively. For ANOVAs, effect size was calculated using partial eta-squared.

In supplementary materials, we also conducted the same analyses but assigned Control A to the Retinotopic Location and Control B to the Spatiotopic Location in Experiment 2 and 3, which was consistent with Shafer-Skelton et al (Shafer-Skelton et al., 2017). Thus, we calculated the spatial congruency bias of spatiotopic and retinotopic conditions by subtracting Same Spatiotopic from Control B and subtracting Same Retinotopic from Control A in Experiment 2 and 3. The results are consistent with the main text reporting the average of Control A and B (Figure S1). Thus, how we choose Control doesn’t affect our conclusions.

Finally, our main analyses focused on the spatial congruency bias, but we also calculated sensitivity (d’) to measure possible facilitation effects. Here, we calculated d’ using signal detection theory: d’ = z(hit rate) – z(false alarm rate). Before we analyzed the data, we excluded trials on which subjects responded with response times (RTs) greater than or less than 2.5 standard deviations from the subject’s mean RT for each subject. Tables S2-9 report the condition mean and statistics for all behavioral measures, including RT, accuracy, d-prime, and proportion ‘Same’ response.

## Results

### Experiments 1 and 2: Dynamic saccade vs static fixation contexts

Experiments 1 and 2 aimed to test whether dynamic saccade context can trigger spatiotopic object-location binding. In Experiment 1 (dynamic saccade context), participants were asked to perform a sequence of 8 eye movements on each trial (Figure 1A). At two points in the sequence, object stimuli appeared on the screen. Participants had to judge whether the two objects were the same or different identity. In Experiment 2 (static context), rather than the sequence of 8 saccades, participants were asked to conduct only one saccade in the middle of the trial, and the two object stimuli were presented during static fixation periods (Figure 1B). Experiment 2 is the same single-saccade Spatial Congruency Bias task that was used in Shafer-Skelton et al., (Shafer-Skelton et al., 2017), but here we designed the timing, stimuli, and experimental design of the trials to perfectly match those in Experiment 1. In other words, Experiment 2 was identical to Experiment 1, save for the dynamic saccade context manipulation.

The task in the two experiments was to judge whether the identities of the two objects were the same or different; the objects could appear in different locations, but location was irrelevant to the task. In both experiments, the critical location conditions were Same Spatiotopic location (objects 1 and 2 appeared in the same absolute screen location) and Same Retinotopic location (objects 1 and 2 appeared in the same eye-centered location), which were compared to Control locations (object 2 appeared in a location that was different from object 1 in both retinotopic and spatiotopic coordinates; Figures 1A-B). To assess object-location binding, we calculated the Spatial Congruency Bias (SCB: Golomb et al. (Golomb et al., 2014)).

The results in Experiment 1 revealed that participants were more likely to report the objects as the same identity when they appeared in the same spatiotopic or same retinotopic locations, compared to the control location (Figure 1G). Paired t-tests revealed a significant spatiotopic SCB (difference in response bias for Control vs. Same Spatiotopic, t(15) = 3.6060, p = 0.0026, d = 0.9366, BF_10_ = 16.937), and a significant retinotopic SCB (Control vs. Same Retinotopic, t(15) = 2.6115, p = 0.0196, d = 0.6571, BF_10_ = 3.152), with no significant difference between spatiotopic and retinotopic biases: t(15) = -0.6686, p = 0.5139, d = 0.1822, BF_10_ = 0.311). This contrasts with the results from Shafer-Skelton et al. (Shafer-Skelton et al., 2017), which only found a significant retinotopic SCB in the more static single-saccade context.

To test whether it was in fact the dynamic saccade context that triggered spatiotopic object-location binding, we conducted Experiment 2 with an otherwise identical design but without the dynamic saccade context. In Experiment 2 (static context), we only found a significant retinotopic SCB (Control vs. Same Retinotopic: t(15) = 8.8710, p < 0.001, d = 2.1290, BF_10_ = 65580), similar to Shafer-Skelton et al (Shafer-Skelton et al., 2017), with no significant spatiotopic SCB (Control vs. Same Spatiotopic: t(15) = 0.4661, p = 0.6478, d = 0.1113, BF_10_ = 0.281) (Figure 1D), and a significantly greater retinotopic than spatiotopic bias (t(15) = -5.5787, p < 0.001, d = 1.5088, BF_10_ = 503).

A 2 × 2 ANOVA on the SCB measures with a within-subjects factor of Reference frame (Spatiotopic or Retinotopic) and an across-subjects factor of Experiment Context (dynamic saccade or static fixation) confirmed a significant interaction between reference frame and experimental context (F(1, 30) = 15.213, p < 0.001, η^2^ = 0.337). The main effect for reference frame was also significant (F(1, 30) = 8.032, p = 0.008, η^2^ = 0.211), while the main effect of experiment context was not (F(1, 30) = 0.170, p = 0.683, η^2^ = 0.006).

These results suggest that object features seem to be bound to an object’s native retinotopic location by default, and remain tied to retinotopic coordinates in more static contexts. But in more dynamic saccade contexts, spatiotopic object-location binding can also develop, suggesting that object representations can be automatically remapped or converted to reflect spatiotopic coordinates. Thus, dynamic saccade context can indeed play a critical role in creating spatiotopic object-location binding. Comparatively, static context only triggered retinotopic object-location binding.

### Experiment 3 and 4: Which factors contribute to dynamic saccade context?

Based on Experiments 1 and 2, we found that static context could only trigger retinotopic object-location binding, but dynamic saccade context could trigger both spatiotopic and retinotopic object-location binding. Design-wise, there were two factors that comprised the dynamic saccade context in Experiment 1: the presence of multiple repeated eye movements, and the fact that stimuli were presented peri-saccadically, with stimulus appearance timed to straddle an eye movement (Figure 2A). To test whether one or both of these factors was critical for triggering spatiotopic object-location binding, we separately manipulated these two factors in Experiments 3 and 4. In Experiment 3 (repeated eye movements only), subjects did 8 sequential saccades on each trial, as in Experiment 1. But here, the stimuli were timed to appear while subjects were stably fixating on one location, as in Experiment 2. In other words, the eye movements occurred before and after, but not during, stimulus presentation in Experiment 3. Contrastingly, in Experiment 4, an eye movement occurred during the stimulus presentation, as in Experiment 1, but here subjects only did 3 saccades on each trial (the minimum necessary for the manipulation; Figure 2A). If being in a dynamic context with multiple repeated eye movements enhances or induces spatiotopic object-location binding, we would expect that Experiment 3 should show a significant spatiotopic SCB, similar to Experiment 1. On the other hand, if the critical aspect of the dynamic context is having eye movements during the stimulus presentation, then we would expect that Experiment 4 should resemble Experiment 1 with a spatiotopic SCB. Finally, if both factors are necessary for spatiotopic object-location binding, we would expect both Experiments 3 and 4 to more closely resemble Experiment 2, with only a retinotopic SCB.

To facilitate comparison across experiments, Figure 2B plots the spatiotopic and retinotopic SCBs for each experiment. Experiment 3 revealed a significant retinotopic SCB (t(15) = 3.8460, p = 0.0016, d = 0.9615, BF_10_ = 25.739), but no significant spatiotopic SCB (t(15) = -1.7728, p = 0.0966, d = -0.4432, BF_10_ = 0.913), and the retinotopic SCB was significantly stronger than the spatiotopic SCB (t(15) = 5.7404, p < 0.0001, d = 1.3849, BF_10_ = 655.613). Experiment 4 also revealed a significant retinotopic SCB (t(15) = 6.0155, p < 0.001, d = 1.5039, BF_10_ = 1024.426). The spatiotopic SCB was not significant (t(15) = 2.0673, p = 0.0564, d = 0.5168, BF_10_ = 1.370), but the Bayes factors suggested inconclusive evidence, with neither strong evidence for the presence or absence of a spatiotopic congruency bias. The retinotopic SCB was significantly greater than the spatiotopic SCB (t(15) = -2.7880, p = 0.0138, d = -0.8844, BF_10_ = 4.201).

The results of Experiments 3 and 4 suggest that both factors – multiple repeated eye movements and eye movements during the stimulus – are important for triggering spatiotopic object-location binding, such that the two dynamic saccade factors may each be necessary, but not individually sufficient, to induce reliable spatiotopic binding. In constrast, retinotopic object-location binding was present in all contexts.

### Across experiments analysis

In order to further compare object-location binding under these different conditions, we conducted an analysis across all four experiments to quantify how the two different factors (repeated eye movements and eye movement during stimulus) influence object-location binding. We first performed a 2 × 2 × 2 ANOVA with a within-subjects factor of Reference frame (Spatiotopic or Retinotopic) and across-subjects factors of Repeated eye movements factor (present or not) and Eye movement during stimulus factor (present or not). A significant main effect was found for reference frame (F(1, 60) = 35.172, p < 0.001, η^2^ = 0.370). The main effects of repeated eye movements factor and eye movement during stimulus factor were not significant (repeated eye movements factor: F(1, 60) = 1.527, p = 0.221, η^2^ = 0.025; eye movement during stimulus factor: F(1, 60) = 3.390, p = 0.0710, η^2^ = 0.053), but there were significant interactions between reference frame and repeated eye movements factor (F(1, 60) = 4.244, p = 0.044, η^2^ = 0.066), and reference frame and eye movement during stimulus factor (F(1, 60) = 12.953, p < 0.001, η^2^ = 0.178). However, the interaction between the two across-subjects factors was not significant (F(1, 60) = 0.859, p = 0.358, η^2^ = 0.014), nor was the 3-way interaction (F(1, 60) = 3.307, p = 0.074, η^2^ = 0.052).

We followed with two separate 2 (Repeated eye movements factor) × 2 (Eye movement during stimulus factor) ANOVAs to assess how these two factors influence spatiotopic and retinotopic object-location binding respectively. For spatiotopic, a significant main effect was found for the eye movement during stimulus factor (F(1, 60) = 13.538, p < 0.001, η^2^ = 0.184), but not for the repeated eye movements factor (F(1, 60) = 0.037, p = 0.848, η^2^ = 0.001). For retinotopic, a significant main effect was found for the repeated eye movements factor (F(1, 60) = 4.616, p = 0.036, η^2^ = 0.071) but not for the eye movement during stimulus factor (F(1, 60) = 0.328, p = 0.569, η^2^ = 0.005). There was no significant interaction between the two factors for either case (spatiotopic: F(1, 60) = 3.445, p = 0.068, η^2^ = 0.054, retinotopic: F(1, 60) = 0.086, p = 0.771, η^2^ = 0.001).

These across-experiments results suggest that both factors influence object-location binding. Having an eye movement during stimulus presentation seems to have a stronger effect on forming more stable spatiotopic object-location binding than the repeated eye movements factor. But as noted earlier, only in the experiment where both factors were present (Experiment 1) was spatiotopic object-location binding reliably triggered. Retinotopic object-location binding, on the other hand, was consistently significant in all four experiments, though the repeated eye movements factor might weaken the magnitude of the effect.

According to our findings from four experiments, there might be a dynamic system of object-location binding. Although the exact mechanism(s) supporting this spatiotopic binding remains to be investigated, we speculate there might be a mechanism whereby an eye movement during the stimulus could be thought of as a gating factor; when present, that triggers the opening of the formation of the spatiotopic representation (but doesn’t influence the existing retinotopic representation). However, opening the gate alone is not enough to reliably trigger spatiotopic binding, and repeated eye movements can thus induce a stronger dynamic brain state that encourages the brain to process the dynamic inputs to maintain visual stability, reinforcing more stable spatiotopic representations, while weakening the retinotopic representations (since repeated eye movements may result in a blurring of retinotopic input if not perfectly aligned).

### Sensitivity effects

The primary focus of this study was to examine the spatial congruency bias measure, which reflects a fundamental influence of object location on object identity judgement, to explore how dynamic saccade context influences object-location binding. However, the design also allows us to examine facilitation effects using the sensitivity measure d-prime. Some previous SCB studies have found a same-location sensitivity effect that co-existed with the spatial congruency bias, while others found only a spatial congruency bias but no sensitivity effect. Table S1 lists the sensitivity effects for the spatiotopic and retinotopic conditions, defined as the difference in sensitivity compared to the control location. The only significant sensitivity effect was retinotopic in Experiment 3. Consistent with the prior studies using the SCB paradigm, the sensitivity measure thus appears to be a less consistent measure, especially compared to the SCB measure, which has proven to be a reliable and theoretically meaningful measure of object-location binding specifically (Cave & Chen, 2017; Golomb et al., 2014; Shafer-Skelton et al., 2017; Starks et al., 2020).

## Discussion

The current study revealed a striking effect with potentially broad theoretical implications: that dynamic saccade context can trigger spatiotopic object-location binding, whereas more static contexts produce retinotopic-only binding. We used a novel experimental design to manipulate the number and timing of eye movements while subjects performed an object identity task, via the Spatial Congruency Bias paradigm (Golomb et al., 2014). For real-world visual stability, it would seem more important to encode an object’s identity with the stable spatiotopic location, instead of the constantly-changing retinotopic location. However, a previous series of studies revealed purely retinotopic object-location binding, for stimuli ranging from gabors to novel objects to faces (Shafer-Skelton et al., 2017). We speculated that a more dynamic saccade context might actually promote a more stable representation across eye movements.

Indeed, when we employed a dynamic saccade context with multiple eye movements during the task, we found both strong spatiotopic and retinotopic spatial congruency biases. In contrast, in Experiment 2 we replicated the finding of only retinotopic object-location binding in a static context similar to Shafer-Skelton et al (2017). Thus, the presence of a more dynamic saccade context can starkly influence the reference frame of object-location binding, allowing spatiotopic effects to robustly emerge. Our results suggest that there can be different reference frameworks of object-location binding under different conditions. This finding is particularly salient because in all conditions, object location was fully task-irrelevant.

Our full set of results suggests that only the most dynamic and ecological experimental condition (Experiment 1) triggered reliable spatiotopic object representations (i.e., a spatiotopic Spatial Congruency Bias). In the partially-dynamic conditions (repeated eye movements factor only or eye movement during stimulus factor only), there was only retinotopic object-location binding. Thus, our study isolated two ecologically key factors -- repeated eye movements and eye movement during stimulus – that can contribute to dynamic saccade context at the behavioral level. To our knowledge, this is the first time these factors have been isolated in this way. For example, a previous study on 3D spatial representations found that experimental blocks in which participants made repeated eye movements while viewing the stimulus induced more tolerant spatial position representations in human visual cortex, compared to experimental blocks containing no eye movements (Zhang et al., 2020). However, multiple factors simultaneously varied between these conditions. Moreover, the eye movements in that study were guided in such a way that overlapping retinotopic visual stimulation could have partially accounted for the results; in contrast, in the current study, the critical eye movement separating the retinotopic and spatiotopic locations was in an orthogonal direction, such that the spatiotopic location had no retinotopic contamination. Another study on spatial localization found that repeated saccades may trigger more optimal integration of spatial updating cues (Poletti et al., 2013), but our current findings reveal that merely increasing the number of eye movements is not enough to flip the reference frame of object-location binding from retinotopic to spatiotopic; stable spatiotopic binding only emerged when there was also the eye movement during stimulus factor. As noted earlier, visual stability involves both stabilization of spatial information across eye movements and integration of spatial information with feature and object representation, and these results could reflect that object-location binding requires additional processes for stable encoding.

The behavioral findings we report here may also provide some insight into and future directions for exploring the neural mechanisms of object-location binding across saccades. In terms of this “hard binding problem” (Cavanagh et al., 2010), a few solutions have been speculated. One way spatiotopic object-location binding could be achieved is through a ‘remapping’ mechanism that updates both spatial location and feature/identity information with each saccade. Spatial remapping is well established (Duhamel et al., 1992; Hartmann et al., 2017; Nakamura & Colby, 2002; Neupane et al., 2016; Poletti et al., 2013; Sommer & Wurtz, 2006; Umeno & Goldberg, 1997, 2001), but feature remapping is still unresolved (Cavanagh et al., 2010; Golomb, 2019; Golomb & Mazer, 2021; Lescroart et al., 2016; Melcher, 2007; O’Herron & von der Heydt, 2013; Subramanian & Colby, 2014). An alternative way to achieve spatiotopic binding is that the binding could be converted to a more stable spatiotopic state, and then would not need to be remapped or rebound with each saccade. We refer to this possibility as ‘spatiotopic state conversion’. While it is possible in principle that this spatiotopic state could overwrite or remove retinotopic location binding altogether, our results suggest that it is more likely this spatiotopic binding would occur in addition to retinotopic representations. However, neural evidence for large-scale spatiotopic organization has also proved controversial (Crespi et al., 2011; D’Avossa et al., 2007; Gardner et al., 2008; Golomb & Kanwisher, 2012; McKyton & Zohary, 2007). Thus, it is difficult to differentiate the specific mechanism of spatiotopic object-location binding through our current study, but our study does suggest that the incorporation of more dynamic saccade context might prove insightful for neural studies of visual stability as well.

In addition, the finding of dynamic saccade context triggering more stable spatiotopic effects may not be specific to object location binding. It is worth exploring whether this principle extends to perception of other visual features, other forms of binding (Treisman, 1996), and other types of visual stability (Bridgeman, 2011), as well as how dynamic saccade context might interact with other cues for visual stability, such as visual landmarks (Deubel, 2004; McConkie & Currie, 1996; Verfaillie, 1997), and working memory. For example, a prior behavioral study explored spatial memory across eye movements and found that free viewing a background scene before the spatial memory stimulus could help facilitate spatiotopic memory (Steinberg et al., 2022); the authors attributed this to the availability of semantic content, but our current findings suggest the possibility that the eye movements themselves could have boosted spatiotopic processing. Another intriguing possibility is that static versus dynamic saccade context may also explain why primarily retinotopic object representations have been found in some neuroimaging studies (Gardner et al., 2008; Golomb & Kanwisher, 2012; Lu et al., 2022).

In sum, our study offers a novel step forward in understanding the complex challenges of visual stability across eye movements and object-location binding. We provide strong evidence for dynamic saccade contexts, including both repeated eye movements and eye movements during stimulus, triggering an integrative brain state that facilitates more stable object-location binding. In the real world, we indeed make eye movements during stimulus and repeated eye movements while observing the world. To some extent, our behavioral results confirm that these two factors are essential components of dynamic saccade context. These findings provide strong evidence that dynamic saccade context can trigger an integrative and dynamic brain state facilitating more stable object-location binding, which is crucial to improved understanding of how the brain achieves visual stability and how we form a stable perception of the dynamic world.

## Acknowledgements

NIH R01-EY025648, NSF 1848939

## Supplementary materials

### Sensitivity effects for all experiments

**Table S1.**
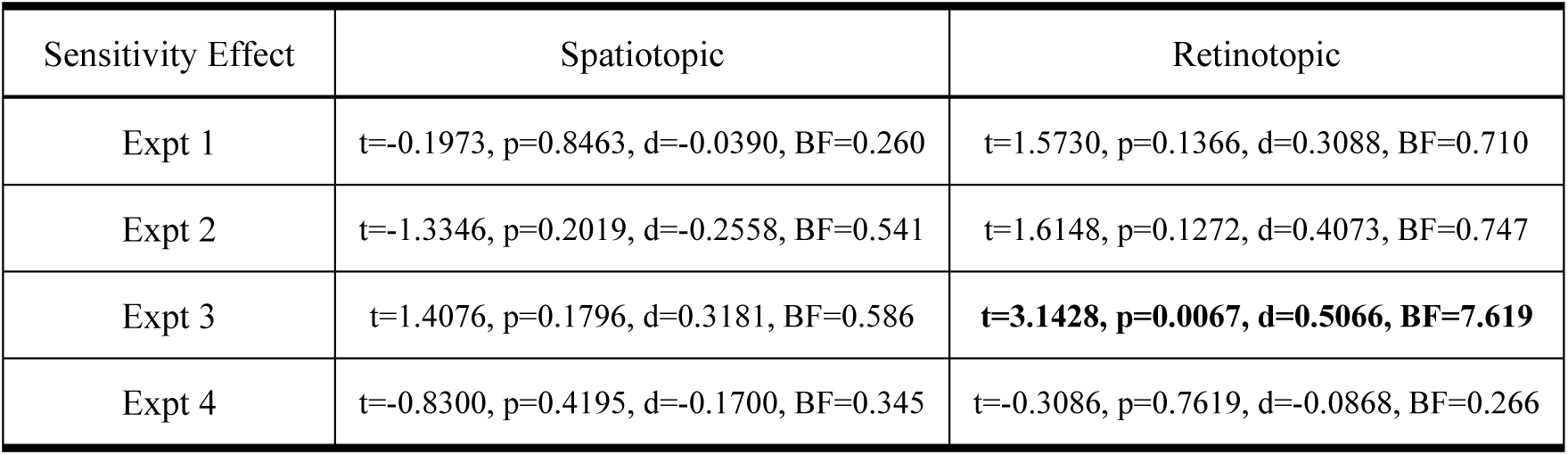
Sensitivity effects for spatiotopic and retinotopic conditions (Spatiotopic sensitivity effect = Same Spatiotopic d-prime minus Control d-prime; Retinotopic sensitivity effect = Same Retinotopic d-prime minus Control d-prime; Control conditions defined as in main text).

| Sensitivity Effect | Spatiotopic | Retinotopic |
| --- | --- | --- |
| Expt 1 | t=-0.1973, p=0.8463, d=-0.0390, BF=0.260 | t=1.5730, p=0.1366, d=0.3088, BF=0.710 |
| Expt 2 | t=-1.3346, p=0.2019, d=-0.2558, BF=0.541 | t=1.6148, p=0.1272, d=0.4073, BF=0.747 |
| Expt 3 | t=1.4076, p=0.1796, d=0.3181, BF=0.586 | **t=3.1428, p=0.0067, d=0.5066, BF=7.619** |
| Expt 4 | t=-0.8300, p=0.4195, d=-0.1700, BF=0.345 | t=-0.3086, p=0.7619, d=-0.0868, BF=0.266 |

### Tables with full results for all behavioral measures and conditions

**Table S2.** Means (and standard deviations) for all behavioral measures and conditions in Experiment 1 (dynamic saccade context condition). SS = Same Spatiotopic; SR = Same Retinotopic; PR = Partial Retinotopic; C = Control

| Expt 1 | Same/Different Identity | SS | SR | PR | C |
| --- | --- | --- | --- | --- | --- |
| RT(s) | Same Identity | 0.6771(0.2741) | 0.6870(0.2935) | 0.6526(0.2875) | 0.6649(0.2752) |
|  | Different Identity | 0.6769(0.2924) | 0.6810(0.2912) | 0.6827(0.2818) | 0.6784(0.3087) |
| Accuracy | Same Identity | 0.7452(0.1244) | 0.7644(0.1069) | 0.7578(0.0988) | 0.6702(0.1203) |
|  | Different Identity | 0.6218(0.1246) | 0.6667(0.1658) | 0.6863(0.1360) | 0.7105(0.1334) |
| P('Same') | Same Identity | 0.7452(0.1243) | 0.7644(0.1069) | 0.7578(0.0988) | 0.6702(0.1203) |
|  | Different Identity | 0.3782(0.1246) | 0.3333(0.1658) | 0.3137(0.1360) | 0.2895(0.1334) |
| d-prime |  | 1.0520(0.5891) | 1.2446(0.5464) | 1.2762(0.6252) | 1.0743(0.5566) |
| Response Bias |  | -0.1916(0.2694) | -0.1362(0.3348) | -0.0934(0.2565) | 0.0679(0.2845) |

**Table S3.**
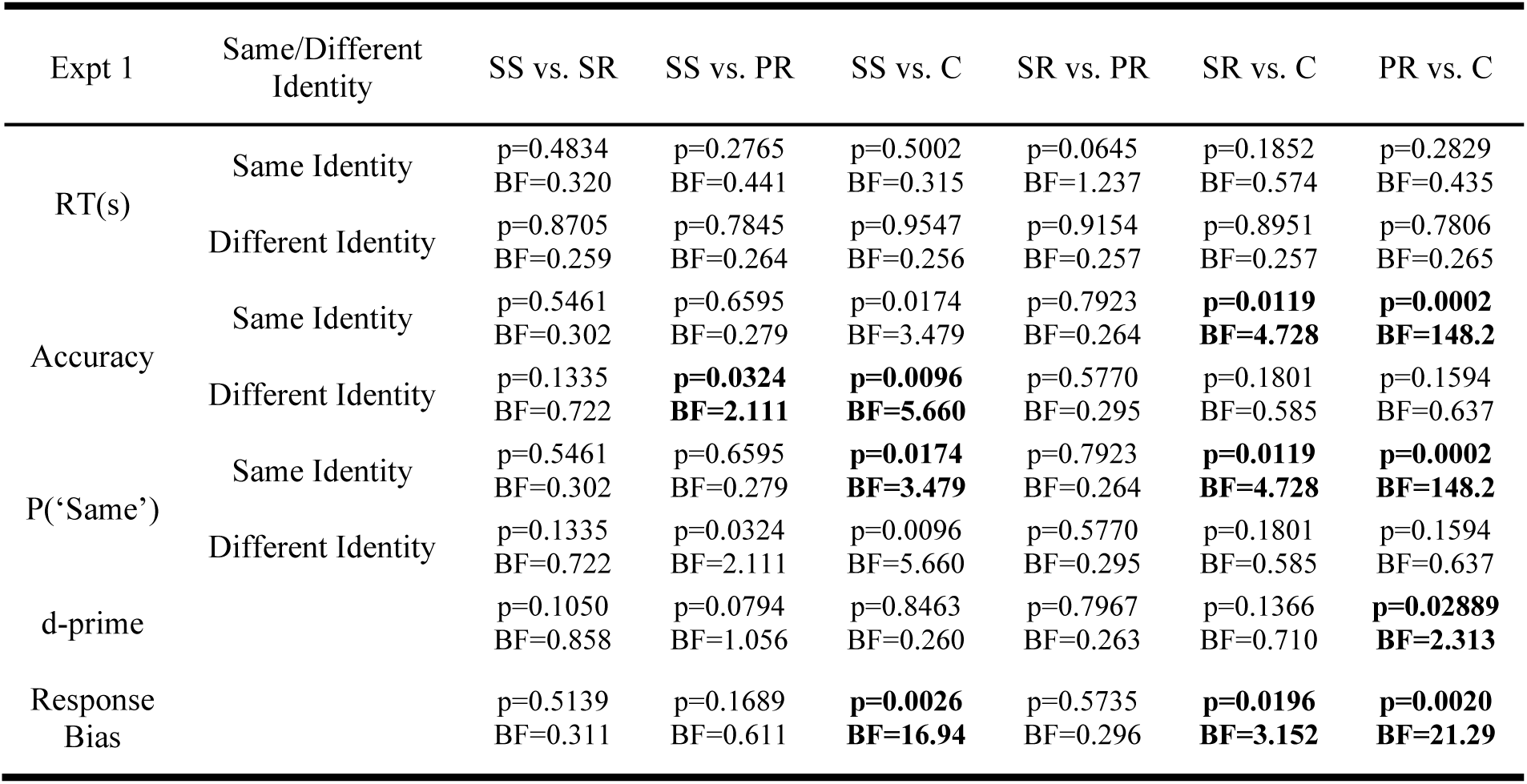
Statistical comparisons, p-values (and BF10), for measures between different location conditions in Experiment 1 (dynamic saccade context condition).

**Table S4.**
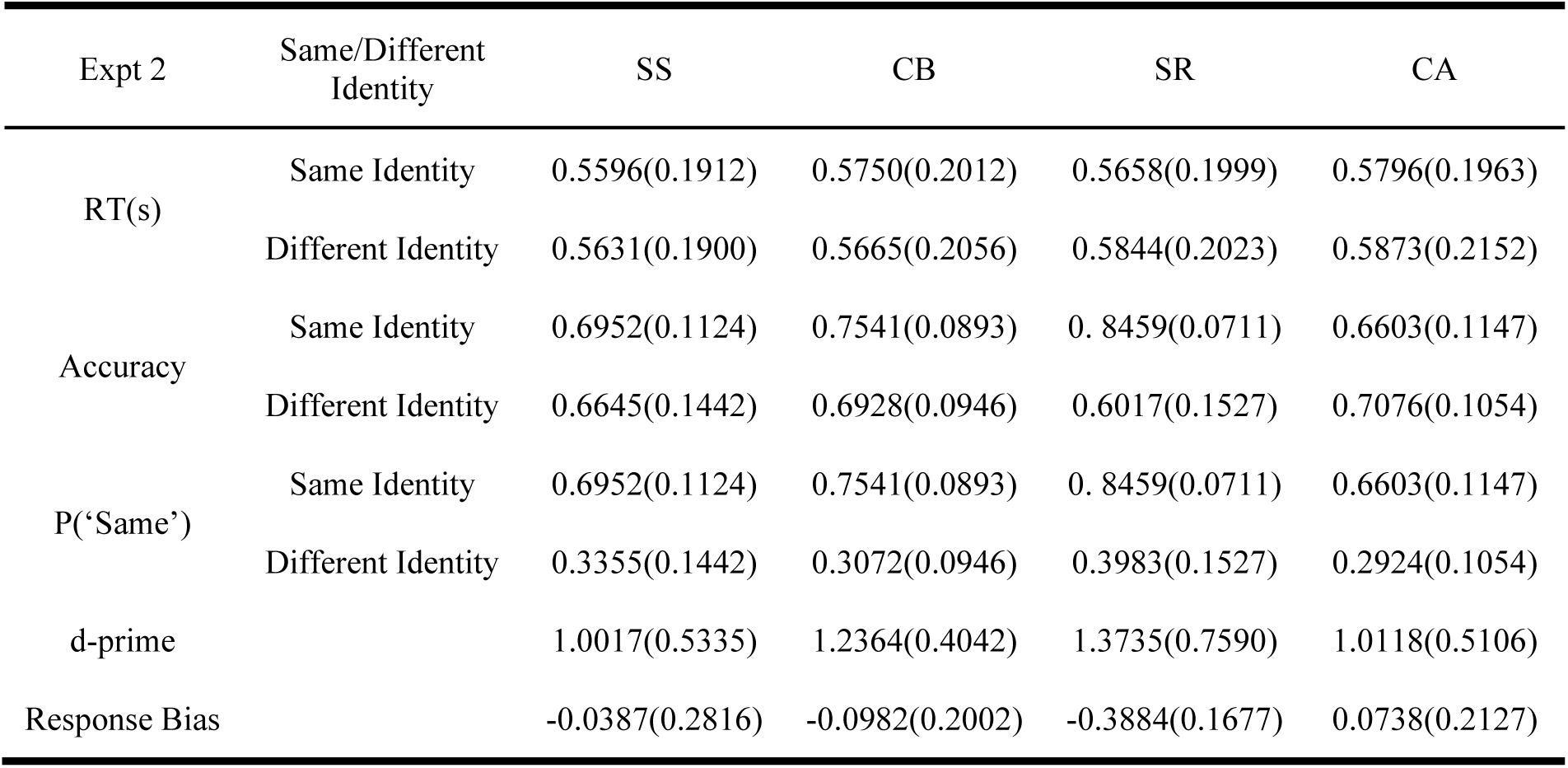
Means (and standard deviations) for all behavioral measures and conditions in Experiment 2 (static context condition). SS = Same Spatiotopic; SR = Same Retinotopic; CA = Control A; CB = Control B.

| Expt 2 | Same/Different Identity | SS | CB | SR | CA |
| --- | --- | --- | --- | --- | --- |
| RT(s) | Same Identity | 0.5596(0.1912) | 0.5750(0.2012) | 0.5658(0.1999) | 0.5796(0.1963) |
|  | Different Identity | 0.5631(0.1900) | 0.5665(0.2056) | 0.5844(0.2023) | 0.5873(0.2152) |
| Accuracy | Same Identity | 0.6952(0.1124) | 0.7541(0.0893) | 0.8459(0.0711) | 0.6603(0.1147) |
|  | Different Identity | 0.6645(0.1442) | 0.6928(0.0946) | 0.6017(0.1527) | 0.7076(0.1054) |
| P('Same') | Same Identity | 0.6952(0.1124) | 0.7541(0.0893) | 0.8459(0.0711) | 0.6603(0.1147) |
|  | Different Identity | 0.3355(0.1442) | 0.3072(0.0946) | 0.3983(0.1527) | 0.2924(0.1054) |
| d-prime |  | 1.0017(0.5335) | 1.2364(0.4042) | 1.3735(0.7590) | 1.0118(0.5106) |
| Response Bias |  | -0.0387(0.2816) | -0.0982(0.2002) | -0.3884(0.1677) | 0.0738(0.2127) |

**Table S5.** Statistical comparisons, p-values (and BF10), for measures between different location conditions in Experiment 2 (static context condition).

| Expt 2 | Same/Different Identity | SS vs. CB | SS vs. SR | SS vs. CA | CB vs. SR | CB vs. CA | SR vs. CA |
| --- | --- | --- | --- | --- | --- | --- | --- |
| RT(s) | Same Identity | p=0.6050<br>BF=0.289 | p=0.7869<br>BF=0.264 | p=0.2363<br>BF=0.488 | p=0.6845<br>BF=0.276 | p=0.8543<br>BF=0.259 | p=0.4498<br>BF=0.332 |
|  | Different Identity | p=0.8953<br>BF=0.257 | p=0.3608<br>BF=0.375 | p=0.4053<br>BF=0.352 | p=0.2702<br>BF=0.447 | p=0.2547<br>BF=0.465 | p=0.8629<br>BF=0.259 |
| Accuracy | Same Identity | **p=0.0132**<br>**BF=4.343** | **p<0.0001**<br>**BF=415.5** | p=0.1026<br>BF=0.873 | **p=0.0003**<br>**BF=115.3** | **p=0.0015**<br>**BF=26.87** | **p<0.0001**<br>**BF=4013** |
|  | Different Identity | p=0.4500<br>BF=0.332 | p=0.0516<br>BF=1.466 | p=0.1788<br>BF=0.588 | **p=0.0283**<br>**BF=2.349** | p=0.4901<br>BF=0.318 | **p=0.0055**<br>**BF=8.926** |
| P('Same') | Same Identity | **p=0.0132**<br>**BF=4.343** | **p<0.0001**<br>**BF=415.5** | p=0.1026<br>BF=0.873 | **p=0.0003**<br>**BF=115.3** | p=0.0015<br>BF=26.87 | **p<0.0001**<br>**BF=4013** |
|  | Different Identity | p=0.4500<br>BF=0.722 | p=0.0516<br>BF=2.111 | p=0.1788<br>BF=5.660 | p=0.5770<br>BF=0.295 | p=0.1801<br>BF=0.585 | p=0.1594<br>BF=0.637 |
| d-prime |  | p=0.0588<br>BF=1.327 | **p=0.0182**<br>**BF=3.351** | p=0.9140<br>BF=0.257 | p=0.3995<br>BF=0.354 | p=0.0367<br>BF=1.915 | **p=0.0456**<br>**BF=1.615** |
| Response Bias |  | p=0.3913<br>BF=0.358 | **p<0.0001**<br>**BF= 502.5** | p=0.0572<br>BF=1.355 | **p<0.0001**<br>**BF=511.4** | **p=0.0019**<br>**BF=22.46** | **p<0.0001**<br>**BF=482600** |

**Table S6.**
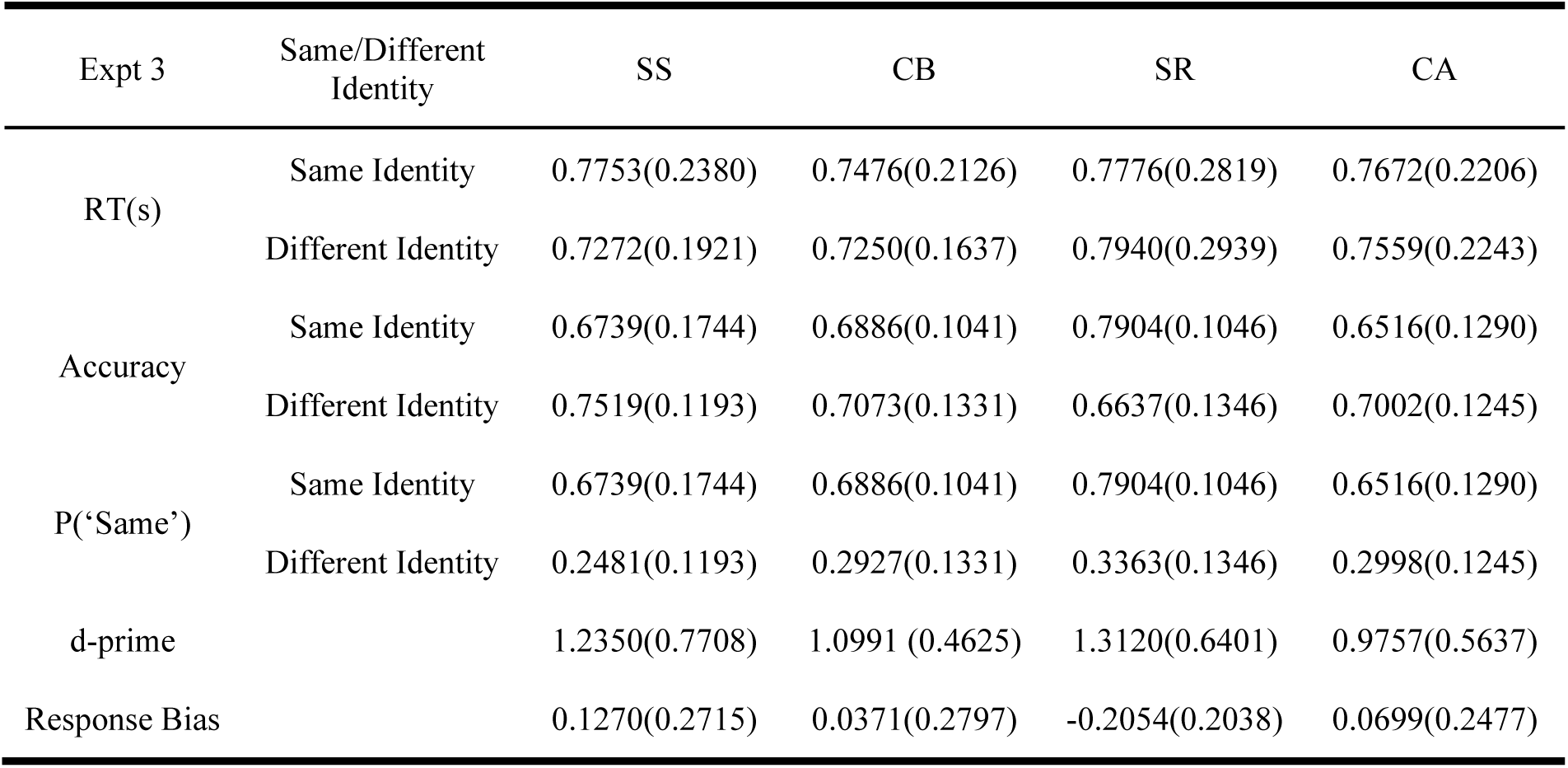
Means (and standard deviations) for all behavioral measures and conditions in Experiment 3 (repeated eye movements only context condition). SS = Same Spatiotopic; SR = Same Retinotopic; CA = Control A; CB = Control B.

| Expt 3 | Same/Different Identity | SS | CB | SR | CA |
| --- | --- | --- | --- | --- | --- |
| RT(s) | Same Identity | 0.7753(0.2380) | 0.7476(0.2126) | 0.7776(0.2819) | 0.7672(0.2206) |
|  | Different Identity | 0.7272(0.1921) | 0.7250(0.1637) | 0.7940(0.2939) | 0.7559(0.2243) |
| Accuracy | Same Identity | 0.6739(0.1744) | 0.6886(0.1041) | 0.7904(0.1046) | 0.6516(0.1290) |
|  | Different Identity | 0.7519(0.1193) | 0.7073(0.1331) | 0.6637(0.1346) | 0.7002(0.1245) |
| P('Same') | Same Identity | 0.6739(0.1744) | 0.6886(0.1041) | 0.7904(0.1046) | 0.6516(0.1290) |
|  | Different Identity | 0.2481(0.1193) | 0.2927(0.1331) | 0.3363(0.1346) | 0.2998(0.1245) |
| d-prime |  | 1.2350(0.7708) | 1.0991 (0.4625) | 1.3120(0.6401) | 0.9757(0.5637) |
| Response Bias |  | 0.1270(0.2715) | 0.0371(0.2797) | -0.2054(0.2038) | 0.0699(0.2477) |

**Table S7.** Statistical comparisons, p-values (and BF10), for measures between different location conditions in Experiment 3 (repeated eye movements only context condition).

| Expt 3 | Same/Different Identity | SS vs. CB | SS vs. SR | SS vs. CA | CB vs. SR | CB vs. CA | SR vs. CA |
| --- | --- | --- | --- | --- | --- | --- | --- |
| RT(s) | Same Identity | p=0.1274<br>BF=0.746 | p=0.9312<br>BF=0.256 | p=0.5291<br>BF=0.307 | p=0.3162<br>BF=0.406 | p=0.1832<br>BF=0.578 | p=0.6994<br>BF=0.274 |
|  | Different Identity | p=0.9098<br>BF=0.257 | p=0.0714<br>BF=1.144 | 0.1374<br>BF=0.707 | p=0.1220<br>BF=0.769 | p=0.2320<br>BF=0.494 | p=0.1932<br>BF=0.558 |
| Accuracy | Same Identity | p=0.7128<br>BF=0.272 | **p=0.0114**<br>**BF=4.896** | 0.3975<br>BF=0.355 | **p=0.0115**<br>**BF=4.858** | p=0.2811<br>BF=0.436 | **p=0.0001**<br>**BF=220.3** |
|  | Different Identity | p=0.1747<br>BF=0.597 | **p=0.0007**<br>**BF=49.06** | p=0.0658<br>BF=1.210 | p=0.1671<br>BF=0.616 | p=0.8142<br>BF=0.262 | p=0.2452<br>BF=0.476 |
| P('Same') | Same Identity | p=0.7128<br>BF=0.272 | **p=0.0114**<br>**BF=4.896** | p=0.3975<br>BF=0.355 | **p=0.0115**<br>**BF=4.858** | p=0.2811<br>BF=0.436 | **p=0.0001**<br>**BF=220.3** |
|  | Different Identity | p=0.1747<br>BF=0.597 | **p=0.0007**<br>**BF=49.06** | p=0.0658<br>BF=1.218 | p=0.1671<br>BF=0.616 | p=0.8142<br>BF=0.262 | p=0.2452<br>BF=0.476 |
| d-prime |  | p=0.4569<br>BF=0.330 | p=0.6276<br>BF=0.285 | **p=0.0775**<br>**BF=1.075** | p=0.1629<br>BF=0.627 | p=0.4182<br>BF=0.346 | **p=0.0003**<br>**BF=106.1** |
| Response Bias |  | p=0.1043<br>BF=0.862 | **p<0.0001**<br>**BF=655.6** | p=0.2572<br>BF=0.462 | **p=0.0039**<br>**BF=11.93** | p=0.5727<br>BF=0.296 | **p=0.0022**<br>**BF=19.15** |

**Table S8.**
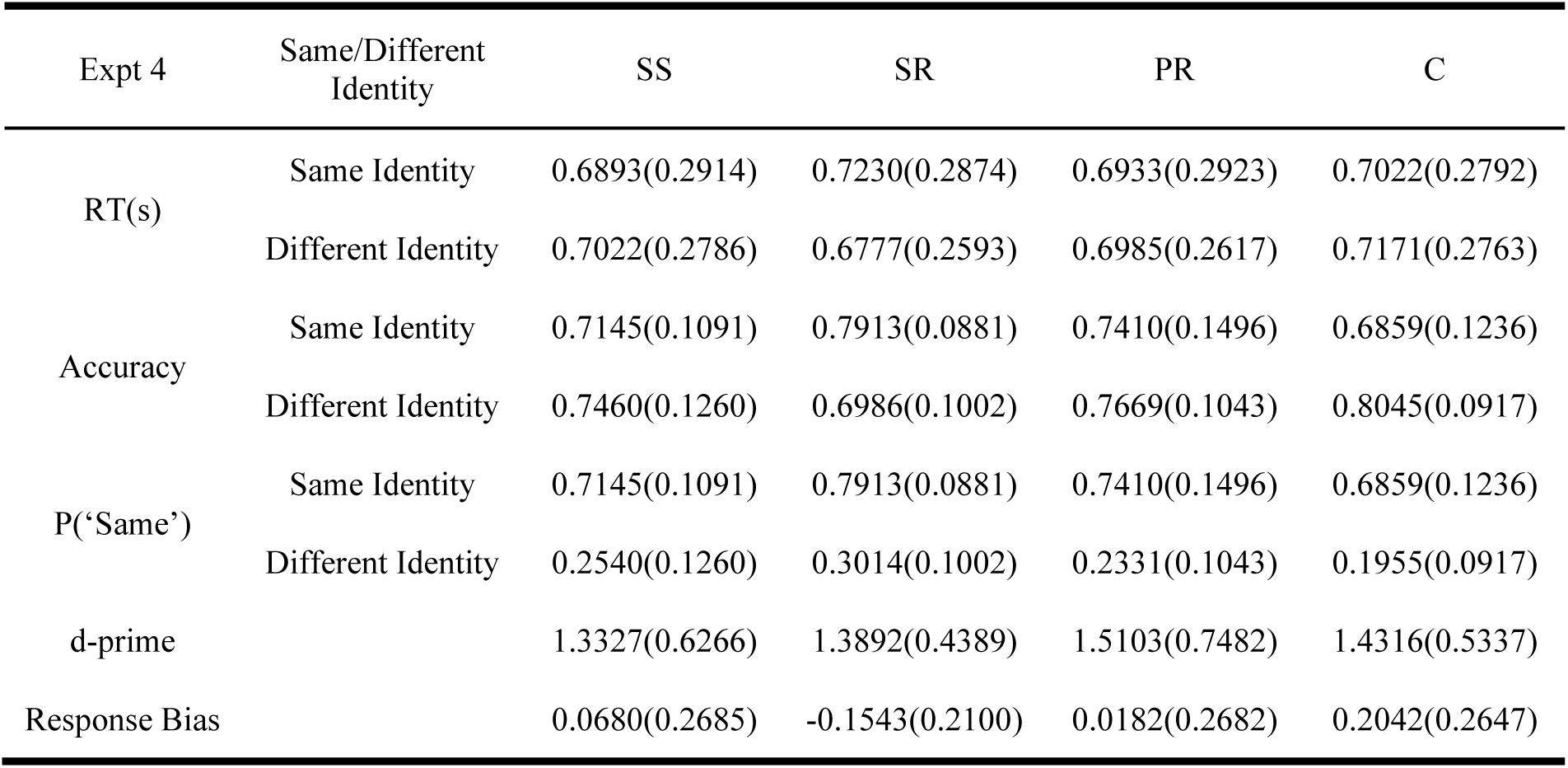
Means (and standard deviations) for all behavioral measures and conditions in Experiment 3 (eye movement during stimulus only context condition). SS = Same Spatiotopic; SR = Same Retinotopic; PR = Partial Retinotopic; C = Control.

| Expt 4 | Same/Different Identity | SS | SR | PR | C |
| --- | --- | --- | --- | --- | --- |
| RT(s) | Same Identity | 0.6893(0.2914) | 0.7230(0.2874) | 0.6933(0.2923) | 0.7022(0.2792) |
|  | Different Identity | 0.7022(0.2786) | 0.6777(0.2593) | 0.6985(0.2617) | 0.7171(0.2763) |
| Accuracy | Same Identity | 0.7145(0.1091) | 0.7913(0.0881) | 0.7410(0.1496) | 0.6859(0.1236) |
|  | Different Identity | 0.7460(0.1260) | 0.6986(0.1002) | 0.7669(0.1043) | 0.8045(0.0917) |
| P('Same') | Same Identity | 0.7145(0.1091) | 0.7913(0.0881) | 0.7410(0.1496) | 0.6859(0.1236) |
|  | Different Identity | 0.2540(0.1260) | 0.3014(0.1002) | 0.2331(0.1043) | 0.1955(0.0917) |
| d-prime |  | 1.3327(0.6266) | 1.3892(0.4389) | 1.5103(0.7482) | 1.4316(0.5337) |
| Response Bias |  | 0.0680(0.2685) | -0.1543(0.2100) | 0.0182(0.2682) | 0.2042(0.2647) |

**Table S9.** Statistical comparisons, p-values (and BF10), for measures between different location conditions in Experiment 4 (eye movement during stimulus context condition).

| Expt 4 | Same/Different Identity | SS vs. SR | SS vs. PR | SS vs. C | SR vs. PR | SR vs. C | PR vs. C |
| --- | --- | --- | --- | --- | --- | --- | --- |
| RT(s) | Same Identity | p=0.1676<br>BF=0.615 | p=0.9072<br>BF=0.257 | p=0.7134<br>BF=0.272 | p=0.3694<br>BF=0.370 | p=0.5414<br>BF=0.303 | p=0.8465<br>BF=0.260 |
|  | Different Identity | p=0.2879<br>BF=0.430 | p=0.9099<br>BF=0.257 | p=0.5936<br>BF=0.291 | p=0.4661<br>BF=0.326 | p=0.0739<br>BF=1.115 | p=0.4914<br>BF=0.318 |
| Accuracy | Same Identity | **p=0.0285**<br>**BF=2.337** | p=0.3206<br>BF=0.403 | p=0.3576<br>BF=0.377 | p=0.1699<br>BF=0.609 | **p=0.0012**<br>**BF=32.92** | **p=0.0414**<br>**BF=1.740** |
|  | Different Identity | p=0.1397<br>BF=0.698 | p=0.3459<br>BF=0.385 | **p=0.0216**<br>**BF=2.917** | **p=0.0308**<br>**BF=2.197** | **p=0.0016**<br>**BF=25.00** | p=0.1277<br>BF=0.745 |
| P('Same') | Same Identity | **p=0.0285**<br>**BF=2.337** | p=0.3206<br>BF=0.403 | p=0.3576<br>BF=0.377 | p=0.1699<br>BF=0.609 | **p=0.0012**<br>**BF=32.92** | **p=0.0414**<br>**BF=1.740** |
|  | Different Identity | p=0.1397<br>BF=0.698 | p=0.3459<br>BF=0.385 | **p=0.0216**<br>**BF=2.917** | **p=0.0308**<br>**BF=2.197** | **p=0.0016**<br>**BF=25.00** | p=0.1277<br>BF=0.745 |
| d-prime |  | p=0.7098<br>BF=0.272 | p=0.1883<br>BF=0.568 | p=0.4195<br>BF=0.345 | p=0.4913<br>BF=0.318 | p=0.7619<br>BF=0.266 | p=0.5265<br>BF=0.307 |
| Response Bias |  | p=0.0138<br>BF=4.201 | p=0.4655<br>BF=0.327 | p=0.0564<br>BF=1.370 | **p=0.0280**<br>**BF=2.372** | **p<0.0001**<br>**BF=1024** | **p=0.0179**<br>**BF=3.395** |

### Analysis with different assignment of control conditions in Experiment 2 and 3

In the main text, we calculated the spatial congruency bias of spatiotopic and retinotopic conditions using a common Control condition baseline. (In Experiments 1 and 4, there was just a single Control location; in Experiments 2 and 3, we averaged Control A and Control B). However, there are other potential ways to assign the control conditions for Experiments 2-3, as discussed in Shafer-Skelton et al (2017). As a supplemental analysis, we tested whether our results would differ if we used a different control assignment. Following Shafer-Skelton et al (2017), we recalculated the spatial congruency bises by subtracting Same Spatiotopic from Control B and subtracting Same Retinotopic from Control A in Experiment 2 and 3. (Spatiotopic and retinotopic spatial congruency biases in Experiment 1 and 4 remain the same.) The pattern of results reported in the main text did not change (see Figure S1, followed by detailed results).

**Figure S1.**
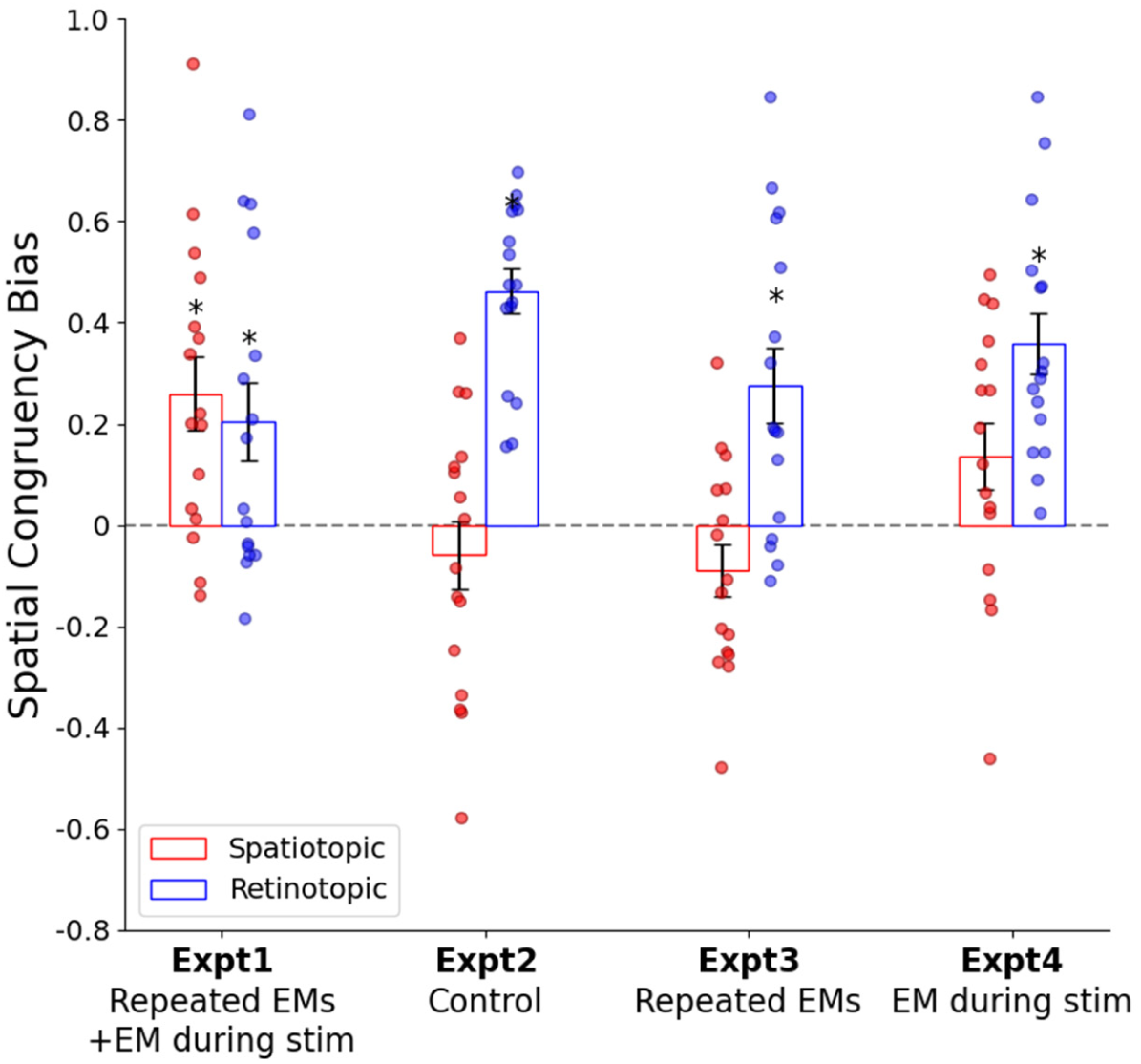
(Compare to main text Figure 2B.) Spatial congruency bias on the identity task re-plotted for spatiotopic and retinotopic conditions based on alternative Control condition assignment (In Experiment 1 and 4: spatiotopic congruency bias: Control bias minus Same Spatiotopic bias; retinotopic congruency bias: Control bias minus Same Retinotopic bias; In Experiment 2 and 3: spatiotopic congruency bias: Control B bias minus Same Spatiotopic bias; retinotopic congruency bias: Control A bias minus Same Retinotopic bias). Error bars are standard error of the mean. Asterisk indicates p<.05 (one-sample t-test for each condition). Spatio = Spatiotopic; Retino = Retinotopic; EM = eye movement, stim = stimulus.

In Experiment 2 (static context), we only found a significant retinotopic SCB (t(15) = 10.4888, p < 0.001, d = 2.4131, BF_10_ = 482600) with no significant spatiotopic SCB (t(15) = - 0.8827, p = 0.3913, d = -0.2433, BF_10_ = 0.358), and a significantly greater retinotopic than spatiotopic bias (t(15) = 5.5787, p < 0.001, d = 1.5088, BF_10_ = 503). In Experiment 3, we also only found a significant retinotopic SCB (t(15) = 3.8460, p = 0.0016, d = 0.9615, BF_10_ = 25.739), but no significant spatiotopic SCB (t(15) = -1.7728, p = 0.0966, d = -0.4432, BF_10_ = 0.913), and the retinotopic congruency bias was significantly stronger than the spatiotopic congruency bias (t(15) = 4.2042, p = 0.0022, d = 1.4864, BF_10_ = 109.615).

The 2 × 2 × 2 ANOVA with a within-subjects factor of Reference frame (Spatiotopic or Retinotopic) and across-subjects factors of Repeated eye movements factor (present or not) and Eye movement during stimulus factor (present or not) revealed a significant main effect for reference frame (F(1, 60) = 40.385, p < 0.001, η^2^ = 0.402). The main effects of repeated eye movements factor and eye movement during stimulus factor were not significant (repeated eye movements factor: F(1, 60) = 1.527, p = 0.221, η^2^ = 0.025; eye movement during stimulus factor: F(1, 60) = 3.390, p = 0.0710, η^2^ = 0.053), but there were significant interactions between reference frame and repeated eye movements factor (F(1, 60) = 6.851, p = 0.0110, η^2^ = 0.102), and reference frame and eye movement during stimulus factor (F(1, 60) = 18.852, p < 0.001, η^2^ = 0.239). However, the interaction between the two factors was not significant (F(1, 60) = 0.859, p = 0.358, η^2^ = 0.014), nor was the 3-way interaction (F(1, 60) = 0.534, p = 0.468, η^2^ = 0.009).

For spatiotopic 2 (Repeated eye movements factor) × 2 (Eye movement during stimulus factor) ANOVA, a significant main effect was found for the eye movement during stimulus factor (F(1, 60) = 17.727, p < 0.001, η^2^ = 0.228), but not for the repeated eye movements factor (F(1, 60) = 0.513, p = 0.476, η^2^ = 0.008). For retinotopic 2 × 2 ANOVA, a significant main effect was found for the repeated eye movements factor (F(1, 60) = 6.767, p = 0.012, η^2^ = 0.101) but not for the eye movement during stimulus factor (F(1, 60) = 1.776, p = 0.188, η^2^ = 0.029). There was no significant interaction between the two factors for either case (spatiotopic congruency bias: F(1, 60) = 1.411, p = 0.240, η^2^ = 0.023, retinotopic congruency bias: F(1, 60) = 0.062, p = 0.8048, η^2^ = 0.001).

## References

Babu, A. S., Scotti, P. S., & Golomb, J. D. (2023). The dominance of spatial information in object indentity judgements: A persistent congruency bias even amidst conflicting statistical regularities. Journal of Experimental Psychology: Human Perception and Performance, 49(5), 672–686.

Bergelt, J., & Hamker, F. H. (2019). Spatial updating of attention across eye movements: A neuro-computational approach. Journal of Vision, 19(7), 10–10.

Brainard, D. H. (1997). The Psychophysics Toolbox. Spatial Vision, 10(4), 433–436.

Bridgeman, B. (2011). Visual stability. The Oxford Handbook of Eye Movements. https://psycnet.apa.org/record/2011-23569-028

Cavanagh, P., Hunt, A. R., Afraz, A., & Rolfs, M. (2010). Visual stability based on remapping of attention pointers. Trends in Cognitive Sciences, 14(4), 147–153.

Cave, K. R., & Chen, Z. (2017). Two kinds of bias in visual comparison illustrate the role of location and holistic/analytic processing differences. *Attention*, Perception, and Psychophysics, 79(8), 2354–2375.

Chen, Z. (2009). Not all features are created equal: Processing asymmetries between location and object features. Vision Research, 49(11), 1481–1491.

Cichy, R. M., Chen, Y., & Haynes, J. D. (2011). Encoding the identity and location of objects in human LOC. NeuroImage, 54(3), 2297–2307.

Crespi, S., Biagi, L., d’Avossa, G., Burr, D. C., Tosetti, M., & Morrone, M. C. (2011). Spatiotopic Coding of BOLD Signal in Human Visual Cortex Depends on Spatial Attention. PLOS ONE, 6(7), e21661.

D’Avossa, G., Tosetti, M., Crespi, S., Biagi, L., Burr, D. C., & Morrone, M. C. (2007). Spatiotopic selectivity of BOLD responses to visual motion in human area MT. Nature Neuroscience, 10, 249–255.

Deubel, H. (2004). Localization of targets across saccades: Role of landmark objects. Visual Cognition, 11(2–3), 173–202.

Duhamel, J. R., Bremmer, F., BenHamed, S., & Graf, W. (1997). Spatial invariance of visual receptive fields in parietal cortex neurons. Nature, 389(6653), 845–848.

Duhamel, J. R., Colby, C. L., & Goldberg, M. E. (1992). The Updating of the Representation of Visual Space in Parietal Cortex by Intended Eye Movements. Science, 255(5040), 90–92.

Gardner, J. L., Merriam, E. P., Movshon, J. A., & Heeger, D. J. (2008). Maps of Visual Space in Human Occipital Cortex Are Retinotopic, Not Spatiotopic. Journal of Neuroscience, 28(15), 3988–3999.

Golomb, J. D. (2019). Remapping locations and features across saccades: a dual-spotlight theory of attentional updating. Current Opinion in Psychology, 29, 211–218.

Golomb, J. D., Albrecht, A. R., Park, S., & Chun, M. M. (2011). Eye Movements Help Link Different Views in Scene-Selective Cortex. Cerebral Cortex, 21(9), 2094–2102.

Golomb, J. D., Chun, M. M., & Mazer, J. A. (2008). The Native Coordinate System of Spatial Attention Is Retinotopic. Journal of Neuroscience, 28(42), 10654–10662.

Golomb, J. D., & Kanwisher, N. (2012). Higher Level Visual Cortex Represents Retinotopic, Not Spatiotopic, Object Location. Cerebral Cortex, 22(12), 2794–2810.

Golomb, J. D., Kupitz, C. N., & Thiemann, C. T. (2014). The influence of object location on identity: A “spatial congruency bias”. Journal of Experimental Psychology: General, 143(6), 2262.

Golomb, J. D., & Mazer, J. A. (2021). Visual Remapping. Annual Review of Vision Science, 7, 257–277.

Goodale, M. A., & Milner, A. D. (1992). Separate visual pathways for perception and action. Trends in Neurosciences, 15(1), 20–25.

Hartmann, T. S., Zirnsak, M., Marquis, M., Hamker, F. H., & Moore, T. (2017). Two types of receptive field dynamics in area v4 at the time of eye movements? Frontiers in Systems Neuroscience, 11, 13.

Kovacs, O., & Harris, I. M. (2019). The role of location in visual feature binding. *Attention*, Perception, and Psychophysics, 81(5), 1551–1563.

Lescroart, M. D., Kanwisher, N., & Golomb, J. D. (2016). No evidence for automatic remapping of stimulus features or location found with fMRI. Frontiers in Systems Neuroscience, 10, 53.

Lu, Z., Shafer-Skelton, A., & Golomb, J. (2022). Gaze-centered spatial representations in human hippocampus. 2022 Conference on Cognitive Computational Neuroscience, 614–616.

Marino, A. C., & Mazer, J. A. (2016). Perisaccadic updating of visual representations and attentional states: Linking behavior and neurophysiology. Frontiers in Systems Neuroscience, 10(FEB), 3.

McConkie, G. W., & Currie, C. B. (1996). Visual Stability Across Saccades while Viewing Complex Pictures. Journal of Experimental Psychology: Human Perception and Performance, 22(3), 563–581.

McKyton, A., & Zohary, E. (2007). Beyond Retinotopic Mapping: The Spatial Representation of Objects in the Human Lateral Occipital Complex. Cerebral Cortex, 17(5), 1164–1172.

Melcher, D. (2007). Predictive remapping of visual features precedes saccadic eye movements. Nature Neuroscience, 10(7), 903–907.

Melcher, D., & Morrone, C. (2003). Spatiotopic temporal integration of visual motion across saccadic eye movements. Nature Neuroscience, 6, 877–881.

Mishkin, M., & Ungerleider, L. G. (1982). Contribution of striate inputs to the visuospatial functions of parieto-preoccipital cortex in monkeys. Behavioural Brain Research, 6(1), 57– 77.

Nakamura, K., & Colby, C. L. (2002). Updating of the visual representation in monkey striate and extrastriate cortex during saccades. Proceedings of the National Academy of Sciences of the United States of America, 99(6), 4026–4031.

Neupane, S., Guitton, D., & Pack, C. C. (2016). Two distinct types of remapping in primate cortical area V4. Nature Communications, 7(1), 1–11.

O’Herron, P., & von der Heydt, R. (2013). Remapping of Border Ownership in the Visual Cortex. Journal of Neuroscience, 33(5), 1964–1974.

Poletti, M., Burr, D. C., & Rucci, M. (2013). Optimal Multimodal Integration in Spatial Localization. Journal of Neuroscience, 33(35), 14259–14268.

Ross, J., & Ma-Wyatt, A. (2003). Saccades actively maintain perceptual continuity. Nature Neuroscience, 7(1), 65–69.

Schwarzlose, R. F., Swisher, J. D., Dang, S., & Kanwisher, N. (2008). The distribution of category and location information across object-selective regions in human visual cortex. Proceedings of the National Academy of Sciences of the United States of America, 105(11), 4447–4452.

Shafer-Skelton, A., Kupitz, C. N., & Golomb, J. D. (2017). Object-location binding across a saccade: A retinotopic spatial congruency bias. *Attention*, Perception, and Psychophysics, 79(3), 765–781.

Snyder, L. H., Grieve, K. L., Brotchie, P., & Andersen, R. A. (1998). Separate body- and world-referenced representations of visual space in parietal cortex. Nature, 394(6696), 887–891.

Sommer, M. A., & Wurtz, R. H. (2006). Influence of the thalamus on spatial visual processing in frontal cortex. Nature, 444(7117), 374–377.

Stanislaw, H., & Todorov, N. (1999). Calculation of signal detection theory measures. Behavior Research Methods, Instruments, & Computers, 31(1), 137–149.

Starks, M. D., Shafer-Skelton, A., Paradiso, M., Martinez, A. M., & Golomb, J. D. (2020). The influence of spatial location on same-different judgments of facial identity and expression. Journal of Experimental Psychology: Human Perception and Performance, 46(12), 1538– 1552.

Steinberg, N. J., Roth, Z. N., & Merriam, E. P. (2022). Spatiotopic and retinotopic memory in the context of natural images. Journal of Vision, 22(4), 11–11.

Subramanian, J., & Colby, C. L. (2014). Shape selectivity and remapping in dorsal stream visual area LIP. Journal of Neurophysiology, 111(3), 613–627.

Sun, L. D., & Goldberg, M. E. (2016). Corollary Discharge and Oculomotor Proprioception: Cortical Mechanisms for Spatially Accurate Vision. Annual Review of Vision Science, 2, 61–84.

Treisman, A. M. (1996). The binding problem. Current Opinion in Neurobiology, 6(2), 171–178.

Treisman, A. M., & Gelade, G. (1980). A feature-integration theory of attention. Cognitive Psychology, 12(1), 97–136.

Treisman, A. M., & Zhang, W. (2006). Location and binding in visual working memory. Memory & Cognition, 34(8), 1704–1719.

Tsal, Y., & Lavie, N. (1988). Attending to color and shape: The special role of location in selective visual processing. Perception & Psychophysics, 44(1), 15–21.

Turi, M., & Burr, D. (2012). Spatiotopic perceptual maps in humans: evidence from motion adaptation. Proceedings of the Royal Society B: Biological Sciences, 279(1740), 3091– 3097.

Umeno, M. M., & Goldberg, M. E. (1997). Spatial processing in the monkey frontal eye field. I. Predictive visual responses. Journal of Neurophysiology, 78(3), 1373–1383.

Umeno, M. M., & Goldberg, M. E. (2001). Spatial processing in the monkey frontal eye field. II. Memory responses. Journal of Neurophysiology, 86(5), 2344–2352.

Verfaillie, K. (1997). Transsaccadic memory for the egocentric and allocentric position of a biological-motion walker. Journal of Experimental Psychology: Learning Memory and Cognition, 23(3), 739–760.

Wexler, M., & Van Boxtel, J. J. A. (2005). Depth perception by the active observer. Trends in Cognitive Sciences, 9(9), 431–438.

Witt, J. K., Taylor, J. E. T., Sugovic, M., & Wixted, J. T. (2015). Signal Detection Measures Cannot Distinguish Perceptual Biases from Response Biases. Perception, 44(3), 289–300.

Zhang, X., Jones, C. M., & Golomb, J. D. (2020). Decoding 3D spatial location across saccades in human visual cortex. BioRxiv, 2020.07.05.188458.

